# Comparison of single gene and module-based methods for modeling gene regulatory networks

**DOI:** 10.1101/307884

**Authors:** Mikel Hernaez, Olivier Gevaert

## Abstract

Gene regulatory networks describe the regulatory relationships among genes, and developing methods for reverse engineering these networks are an ongoing challenge in computational biology. The majority of the initially proposed methods for gene regulatory network discovery create a network of genes and then mine it in order to uncover previously unknown regulatory processes. More recent approaches have focused on inferring modules of co-regulated genes, linking these modules with regulator genes and then mining them to discover new molecular biology.

In this work we analyze module-based network approaches to build gene regulatory networks, and compare their performance to the well-established single gene network approaches. In particular, we focus on the problem of linking genes with known regulatory genes. First, modules are created iteratively using a regression approach that links co-expressed genes with few regulatory genes. After the modules are built, we create bipartite graphs to identify a set of target genes for each regulatory gene. We analyze several methods for uncovering these modules and show that a variational Bayes approach achieves significant improvement with respect to previously used methods for module creation on both simulated and real data. We also perform a topological and gene set enrichment analysis and compare several module-based approaches to single gene network approaches where a graph is built from the gene expression profiles without clustering genes in modules. We show that the module-based approach with variational Bayes outperforms all other methods and creates regulatory networks with a significantly higher rate of enriched molecular pathways.

The code is written in R and can be downloaded from https://github.com/mikelhernaez/linker.

## Introduction

Reverse engineering gene regulatory networks from gene expression data is still a major challenge in computational biology (Gevaert et al. 2007). Gene regulatory networks define regulatory relationships among genes including for example relationships between transcription factors and their target genes. They provide a concise representation of the transcriptional regulatory landscape of the cell, and although they do not reflect post-transcriptional modifications, gene regulatory networks have been successfully used in many applications to elucidate new biological mechanisms and gene-level relationships in cells (Margolin et al. 2006; Akavia et al. 2010; Logsdon et al. 2015).

Most initially proposed methods generate gene regulatory networks by creating a single graph, where each of the nodes is a gene, and an edge between two genes (i.e., nodes) is drawn if a significant relationship between both genes is found. Hence, the key to building highly informative gene regulatory networks is to find the most informative function that provides an accurate measure of the gene-gene relationships such as correlation (Langfelder and Horvath 2008), mutual information (Margolin et al. 2006), and regression models like LASSO (Friedman et al. 2008) and variational Bayes (Logsdon et al. 2015).

Once the graph is created, the next step is usually to look for “hub” genes (i.e. highly connected nodes in the network), or for subnetworks of highly connected genes conveying some biological meaning. In recent years, several of these methods have been successfully applied to uncover new major gene hubs (Basso et al. 2005), discover new molecular targets (Horvath et al. 2006)), or analyze master regulator proteins in tumor samples (Califano and Alvarez 2017).

An alternative approach to the one mentioned above is to directly compute modules of co-expressed genes from the initial gene expression profiles. Some examples of this approach are CONEXIC (Akavia et al. 2010), AMARETTO (Champion et al. 2018; Gevaert et al. 2013; Gevaert and Plevritis 2013) and CaMoDi (Manolakos et al. 2014). Although the module-based approach has not been as well studied as the single gene network approach, it has been used to successfully discover dependencies between known driver genes and tumor dependencies (Champion et al. 2018; Akavia et al. 2010)), or discover new genes for Crohn’s disease (Karczewski et al. 2014).

Several of the aforementioned methods were proposed for relatively small microarray datasets and do not scale well to bigger datasets (see (Yu et al. 2017) and references therein). RNA-sequencing technology has now made genome-wide expression profiling projects on large-scale populations feasible. For example, pan-cancer projects like the PanCanAtlas of The Cancer Genome Atlas (TCGA) (Hoadley et al. 2018; Malta et al. 2018; Campbell et al. 2018) have generated genome wide multi-omic data across more than 10,000 patients spanning a total of 33 tumor types.

In this work we propose a novel algorithm drawing from the module-based methods. Specifically, we first find modules of co-regulated genes whose expression profile can be explained by a combination of few regulatory or driver genes. We analyze several methods for uncovering these modules and show that a variational Bayes approach, used previously in the context of genome wide association studies, achieves significant improvement with respect to the previously used methods for module creation on both simulated and TCGA data. After the modules are built we create bipartite graphs from each of them, yielding highly informative and concise gene regulatory networks. This whole process is bootstrapped several times using a different set of samples and a different random initialization. Finally, using data from the TCGA project we perform a thorough network topology assessment and gene set enrichment analysis of our proposed method, and compare it to single gene network approaches for modeling regulatory networks. We show that the proposed approach generates highly informative networks which contain a significantly higher rate of different enriched elements than the non-module approach.

## Methods

The method described hereafter get as input two gene expression matrices. Namely, a matrix 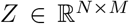, where each of its *N* rows is the *M*-sample-long expression profile of a given gene; and analogously, the matrix 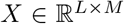 containing the expression profile of *L* regulatory genes. These expression profiles are standardized to have zero mean and standard deviation one.

### Overview of the proposed method

The aim of the proposed method is to find relatively small networks that link few regulatory (or driver) genes with a similarly-regulated set of genes, also known as the *target genes*.

In order to build such networks, the method is divided into two phases. During Phase I the method generates *K* modules of similarly-expressed genes and then associates each module to few regulators (Figure 1). Due to the non-convex nature of the problem, we perform *B* bootstraps of this step with random initializations in order to explore more broadly the set of potentially valid modules (section*). Thus, at the end of this step the method has generated *K* · *B* modules of similarly regulated genes, each of them with their associated regulators.

**Figure 1:**
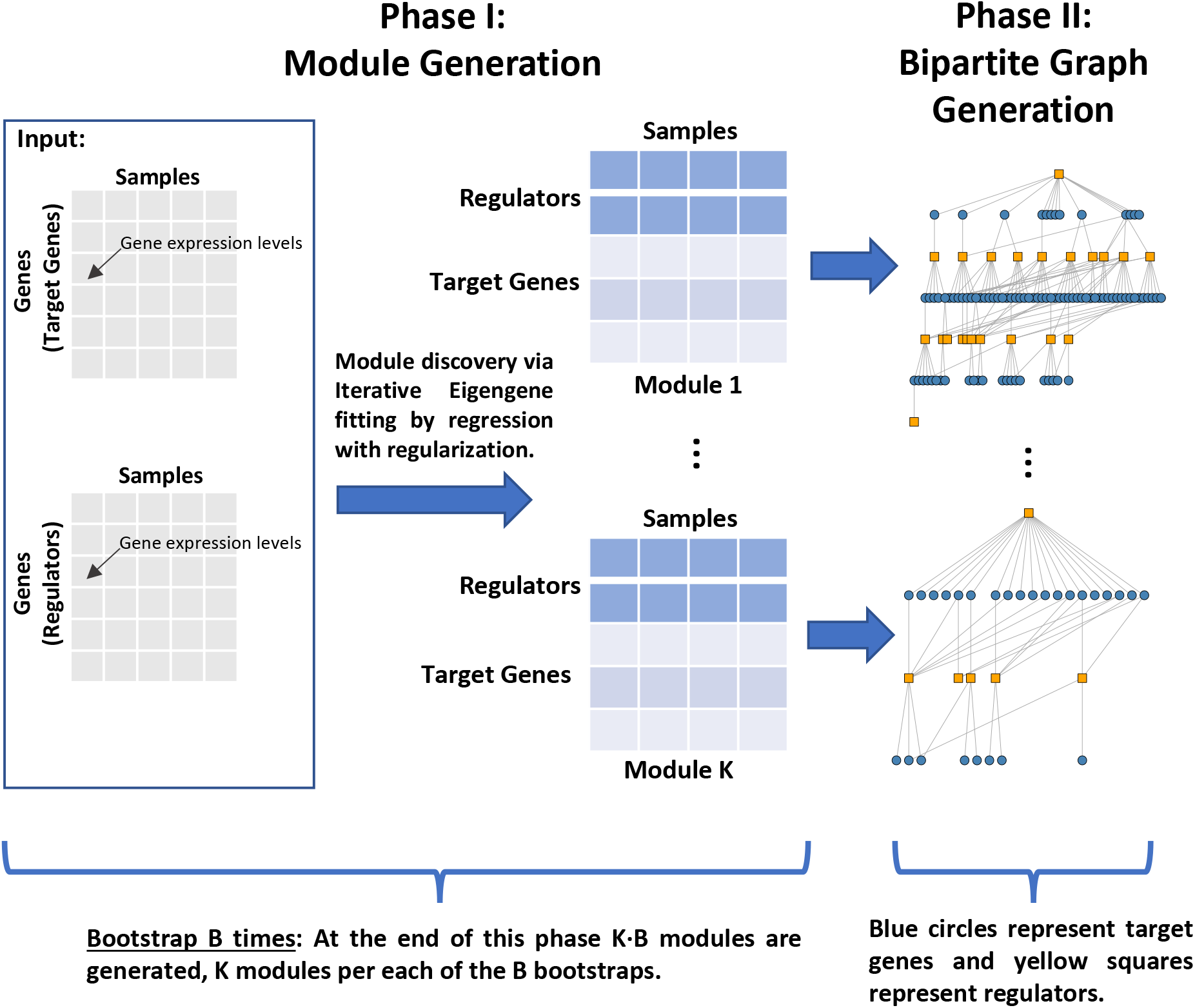
Schematic of the proposed method: Phase I generates several plausible modules from the original data. Phase II generates several bipartite graphs out of the modules generated during Phase I. These bipartite graphs connect the regulators and target genes and are well suited for further analysis and interpretation.

Note that several of the generated *K* · *B* modules will contain genes that were constantly clustered together across bootstraps; thus, those modules will be highly similar to each other. On the other hand, there will be other modules that are unique across the bootstraps, since one of the bootstraps could have ended in a local minima that was not found by any of the other bootstraps. Furthermore, there could be also several modules whose genes were clustered together due to the heuristic nature of the method rather than an underlying regulator-target gene relationship.

If each bootstrap was to generate very similar solutions, the resulting *K* · *B* modules would be composed by *K* distinct set of modules, each containing *B* highly similar modules. (Manolakos et al. 2014) made an argument that such scenario is desirable, however, we believe that such scenario means that the method is not exploring the whole set of possible solutions, and hence not ideal.

At this point, the proposed method has effectively reduced the original problem from having one large set of target genes and another large set of unassociated regulators to having *K* · *B* significantly smaller sets of target genes with their associated regulators.

During Phase II (Figure 1 (right)) the proposed method generates, for each module, a bipartite graph that links the individual target genes to their associated regulators (section*). Note that if no combination of regulators represents accurately the expression profile of a given target gene, that gene is removed from the graph. This scenario arises when the target gene in question was an outlier on the corresponding module.

The output of the proposed method is therefore a set of bipartite graphs that exhibit strong regulator-target gene relationships. These type of graphs are well suited for further analysis and interpretation of the existing regulator-target genes relationships.

In what follows we describe the two phases in more detail.

### Phase I: Module generation

The goal of this step is to generate *K* modules of similarly expressed target geness and relate them to a linear combination of very few regulators.

Broadly described, this step works similarly to the k++ clustering algorithm (Arthur and Vassilvitskii 2007). It first starts with a tailored initialization of the module representatives (as described below). After selecting the most suitable representative for each module, it assigns each gene to the module whose representative is most correlated with in absolute value. After the first cluster assignment is concluded, the *eigengene* (Langfelder and Horvath 2007) of each module is computed (see below).

Each eigengene is then approximated by a linear combination of very few regulators. These approximations become the new representatives and each target gene is then re-assigned to the module that contains the most correlated representative. This process runs iteratively until very few reassignments are made, or until a finite number of iterations are run. Furthermore, we perform *B* bootstraps of this process such that at its conclusion it generates *K* · *B* different modules with their associated regulators. In each bootstrap we select at random 80% of the samples and the initialization of the algorithm is also randomly initialized (Figure 2). In what follows we explain the initialization, the eigengene computation and the approximation by regulators in more detail.

**Figure 2:**
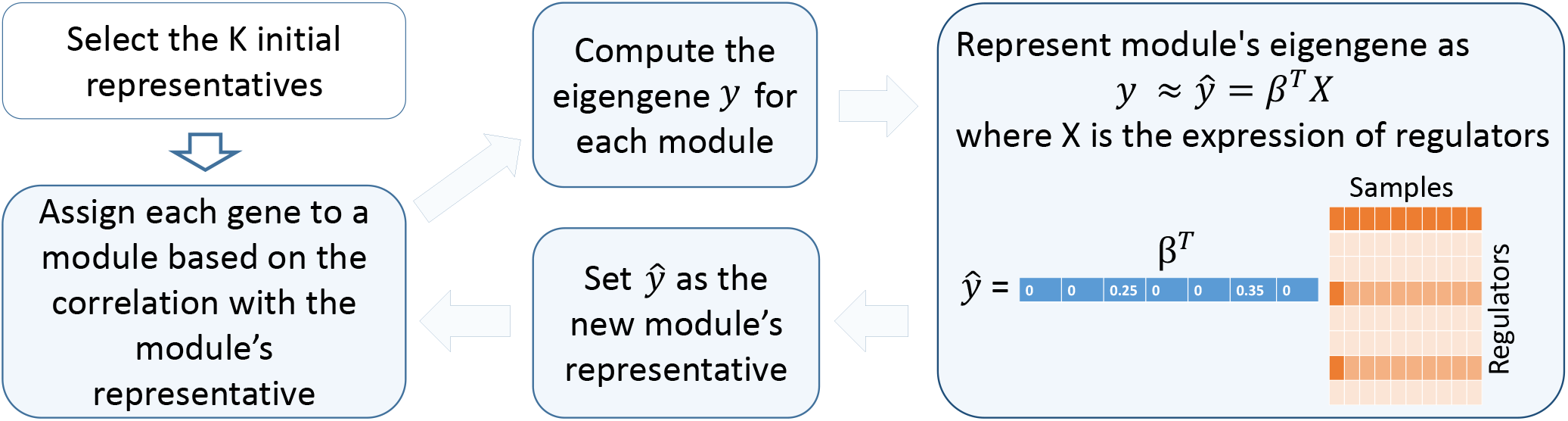
Schematic of the process for generating the different modules during Phase I of the proposed method. This process is bootstrapped *B* times selecting in each bootstrap a different set of samples and a different initialization.

#### Initialization

Phase I takes as input the target gene expression matrix *Z* and the regulator expression matrix *X*. In order to facilitate the convergence of the proposed method we initialize the method as following. First, a target gene is selected at random as the first representative. Then, the method looks for another target gene whose expression is as orthogonal as possible to that of the firstly chosen representative.

The reminding ones are selected as follows. For each of the target genes, a probability proportional to the minimum among the inner products between that gene and the already computed representatives is assigned. That is, a target gene with very low minimum correlation will get a high probability assigned. The next representative is chosen by sampling from the newly computed distribution. This process is performed until all *K* representatives are chosen.

#### Eigengene computation

Once the initial set of modules is built, a representative of each module needs to be selected. For example, in the well-known k-means algorithm (Friedman et al. 2001) the representative of each of the clusters is the centroid, that is, the mean of all the components of the cluster. This approach was also followed by AMARETTO (Gevaert et al. 2013) and CaMoDi (Manolakos et al. 2014). However, we believe that it may not be the most appropriate approach as it forces to ignore negative correlations, since otherwise the mean could be close to zero. Thus, the proposed method choses as the representative of each of the modules the expression profile *y* that is most correlated, on average, with all the components of the cluster, that is, the first eigenvector (i.e., the eigenvector associated with the largest eigenvalue) of the sample covariance matrix of the cluster. Specifically, and with some abuse of notation, a given module 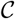 is composed of the different expression profiles 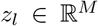, with *M* being the number of samples. The expression matrix *Z* is the matrix whose rows are the expression profiles *z*_*l*_. Thus, the representative of the cluster is the first eigenvector of the sample covariance matrix *Z^T^ Z*. Moreover, the quantity 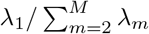 is the percentage of variance in the cluster that is explained by the representative, where *λ*_*m*_ is the eigenvalue associated with the *m*th eigenvector. It can be shown that no other expression profile explains more variation that the first eigenvector (Friedman et al. 2001).

#### Eigengene representation using regulators

This process tries to link the expression of each eigengene *y* with the expression of few regulators. Hence, the aim of this step is to find a vector of weights *β* such that a linear combination of few regulators is as close as possible to the eigengene *y*. We further denote 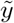 to be the result of this linear combination; hence, 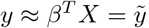.

In this work we test four different ways of estimating *β*. In particular, we use:

- <u>LASSO</u>: It is the method of choice of some of the previously proposed methods like AMARETTO (Gevaert et al. 2013; Champion et al. 2018) or CaMoDi (Manolakos et al. 2014). In order to compute *β* it solves the following optimization problem:

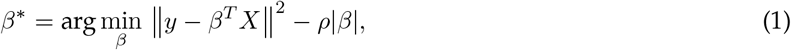

where the value of *ρ* is chosen via 10-fold cross-validation. Specifically, we choose two different solutions of the optimization problem, namely *β_min_* and *β*_1*se*_. The former is computed by selecting the *λ* that minimizes the cross-validation error, while the latter is computed by selecting the sparsest solution such that the cross-validation error is within one standard error from the minimal one. In the following we denote this methods as LASSO_*min*_ and LASSO_1*se*_.
- <u>VBSR</u>: This method was proposed in (Logsdon et al. 2012) and first used to link known regulator genes to the expression of protein-coding genes in (Logsdon et al. 2015). VBSR assumes that *β* is a random variable whose components *β*_*i*_ are distributed as

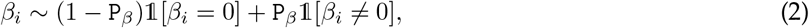

where P_*β*_ is a hyper-parameter determining the prior probability that the component *i* has non-zero probability. To estimate the posterior distribution of the regression coefficients 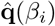, VBSR uses a variational Bayes approximation and minimizes the Kullback-Leibler divergence between the factorized approximate posterior distribution and the full posterior distribution. It was shown in (Logsdon et al. 2012) that the approximate variational Bayes posterior distribution for an arbitrary coefficient *β*_*i*_ is given by

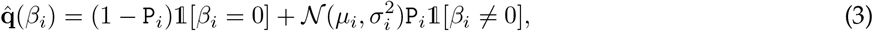

where P_*i*_ is the approximate posterior probability of *β*_*i*_ not being zero, *μ*_*i*_ is the approximate posterior mean of the non-zero effect and 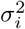 of is the approximate posterior variance of the non-zero effect. The exact expression of these statistics can be found in (Logsdon et al. 2012, Supp. Data). The posterior probability of a coefficient *β_i_* bounds the influence of the variable within the model, e.g., if the posterior probability is zero, then the associated regulator will have no effect on the eigengene in question, and if the posterior probability is one, then the regulator will have a full unpenalized effect. After fitting the model, that is, computing the regression components *β*_*i*_, VBSR defines the Z-statistic for the *i*th component as *z_i_ = μ_i_*/*σ*_*i*_. The Z-statistic is a measure of the significance of the *β*_*i*_ parameter given the rest of the fitted model. Finally, the prior hyper-parameter P_*β*_ is chosen such that the approximate posterior for any *β*_*i*_ is calibrated to 0.95 if its associated Z-statistic passes a Bonferroni correction across the regulators of all models of a family-wise error rate of 0.05. Hence, the *β*_*i*_s associated with a *z*_*i*_ larger than 0.05/(*M* * *N*) are set to zero and the method assumes that there is no mutual effects between the regulator and the eigengene in question.
- <u>LM</u>: In this work we test a method for building correlation networks (or analogously linking eigengenes to regulators) similar to the one used in (Logsdon et al. 2015) and denote it LM. Using correlation between expression levels of genes to build gene regulatory networks has been well studied with considerable success, being WGCNA (Langfelder and Horvath 2008) one of the most used methods. In particular, we fit an ordinary linear model between the eigengene and each of the potential regulators; that is, for each regulator *x*_*i*_ we approximate the eigengene *y* as *y* ≈ *β*_*i*_*x*_*i*_, where *β*_*i*_ is obtained by ordinary least squares. Then, we test the significance of each *β*_*i*_ using the standard Wald test to compute the t-statistic *t*_*i*_. Since we need to set a hard threshold to determine if a relationship between the eigengene *y* and the regulator *x*_*i*_ exists, we apply a Bonferroni correction for *N*^(*j*)^ × *L*^(*j*)^ tests to each pairwise test, where *N*^(*j*)^ is the number of target genes and *L*^(*j*)^ the number of regulators on module *j*. Finally, if the p-value associated to the t-statistic of the pairwise test is smaller than 0.05 after the Bonferroni correction, we consider that a significant relation exists between *y* and *x*_*i*_.

### Phase II: Bipartite graphs generation

The newly generated modules are still difficult to interpret and analyze; hence, during the final phase of the proposed method, each of the modules are converted into a set of bipartite graphs. A bipartite graph is a set of graph nodes decomposed into two disjoint sets such that no two graph nodes within the same set are adjacent.

Figure 1 (right), shows an example of a bipartite graph, in which the square nodes represent different regulators and the circle nodes target genes. An edge is drawn between a square node an a circle node, if the expression profile of the target gene can be partially explained by the expression profile of the regulator. We test the same methods described in section* to find set of regulators within the module that best characterize the expression profile of the target gene at hand.

### Single gene network approach

Phase II could be directly applied to the initial expression matrices yielding a single bipartite graph. Building single gene networks from the input gene expression data has been widely used in the literature and it still reminds the most common approach (see (van Dam et al. 2017) for a review).

Hence, we generate single bipartite graphs from the initial expression matrices using the same methods as in the module-based approach. Thus, for each target gene in the input gene expression matrix we apply either LASSO_*min*_, LASSO_1*se*_, VBSR or LM as described in section* to find which regulators are associated with the target gene. Then we draw an edge in the graph if the corresponding method considers that a relationship exists.

## Results and Discussion

We first describe the data upon which we will evaluate the different approaches discussed above. Specifically, we use the gene expression datasets from The Cancer Genome Atlas (TCGA) project, namely, the ovarian cancer (TCGA-OV) and the head and neck squamous carcinoma (TCGA-HNSC) datasets. Each of these datasets contain the expression values of more than 20,000 genes. However, a significant fraction of these genes are either not expressed in any of the samples or their expression values are not informative. Hence we retain only the 10,000 most varying (i.e., informative) genes. This is also done to considerably reduce the computation time and memory requirements of the tested methods. The regulatory genes are a set of genes, identified via certain biological regulatory mechanisms, which are known to drive other genes. This set has been created based on transcription factor data extracted from the HPRD database (Vaquerizas et al. 2009). Our dataset consists of 3609 regulatory genes.

### Performance analysis on simulated data

Since most of the workload of the proposed method occurs on Phase I, we start by performing a thorough assessment of the different used methods for linking the eigengenes to the regulators.To this end, we simulate expression data such that the true underlying modules and their relationships with the regulators are known.

Let us assume that the expression data *Z* of the target genes is formed by *K* true modules and that the eigengene of each module *j =* {1, …, *K*} is given by *β*^(*j*)^*T*^^ *X*, where *X* is the expression profile matrix of the regulators, taken from the TCGA-OV dataset, and *β*^(*j*)^ is the set of weights that determines the effect that each regulator has on the expression profile of the eigengene. Hence, the expression profile of a target gene *l* from module *j* is given by 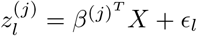, with *l* = {*1*, …, *N_j_*}, *N_j_* the number of genes associated to module *j*, and 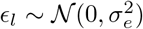 being the biological noise.

Since we expect very few regulators to have a meaningful effect on the eigengene, we generate *β*^(*j*)^ as follows. Let us denote n the number of non-zero elements of *β*^(*j*)^, then we set 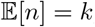, that is, on average, only *k* target genes will have their associated weight 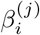 to be non-zero. Furthermore, the non-zero weights are generated as 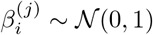.

Due to the non-sparse nature of LM, we omit it from the analysis as, with the set threshold, it was representing each eigengene using significantly more regulators than the other methods (and the true expected value). Moreover, in order to make the comparison as fair as possible, we set the maximum number of regulators used by LASSO to be similar to VBSR. This caused LASSO_1*se*_ and LASSO_*min*_ to perform almost identically. Hence, for the simulated data analysis we compare LASSO to VBSR and analyze their capabilities to recover the true underlaying modules and connections.

#### Regulator-target gene relationship estimation

We first assume that the module assignments are known, that is, we know beforehand to which module *j* each target gene 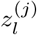 belongs to. This is equivalent to estimating *β*^(*j*)^ given the expression profiles of the target genes belonging to the corresponding module *j*.

Figure 3 shows the estimation performance of LASSO (left) and VBSR (right) for a noise level 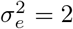. Every blue dot in this figure is a 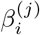 for all *K* = 100 clusters. This shows that VBSR calls significantly fewer false positives with similar number of true positives as LASSO. We define a true positive when both the real and estimated 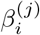 are nonzero, and a false positive when a non-zero effect was estimated when there was no effect. True negatives and false negatives are computed analogously. Furthermore, VBSR estimates the specific effect of each regulator (i.e., the actual values of 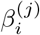) more accurately than LASSO (MSE_LASSO_ = 0.18 vs. MSE_VBSR_ = 0.13, where the MSE is computed as the mean of the differences squared).

**Figure 3:**
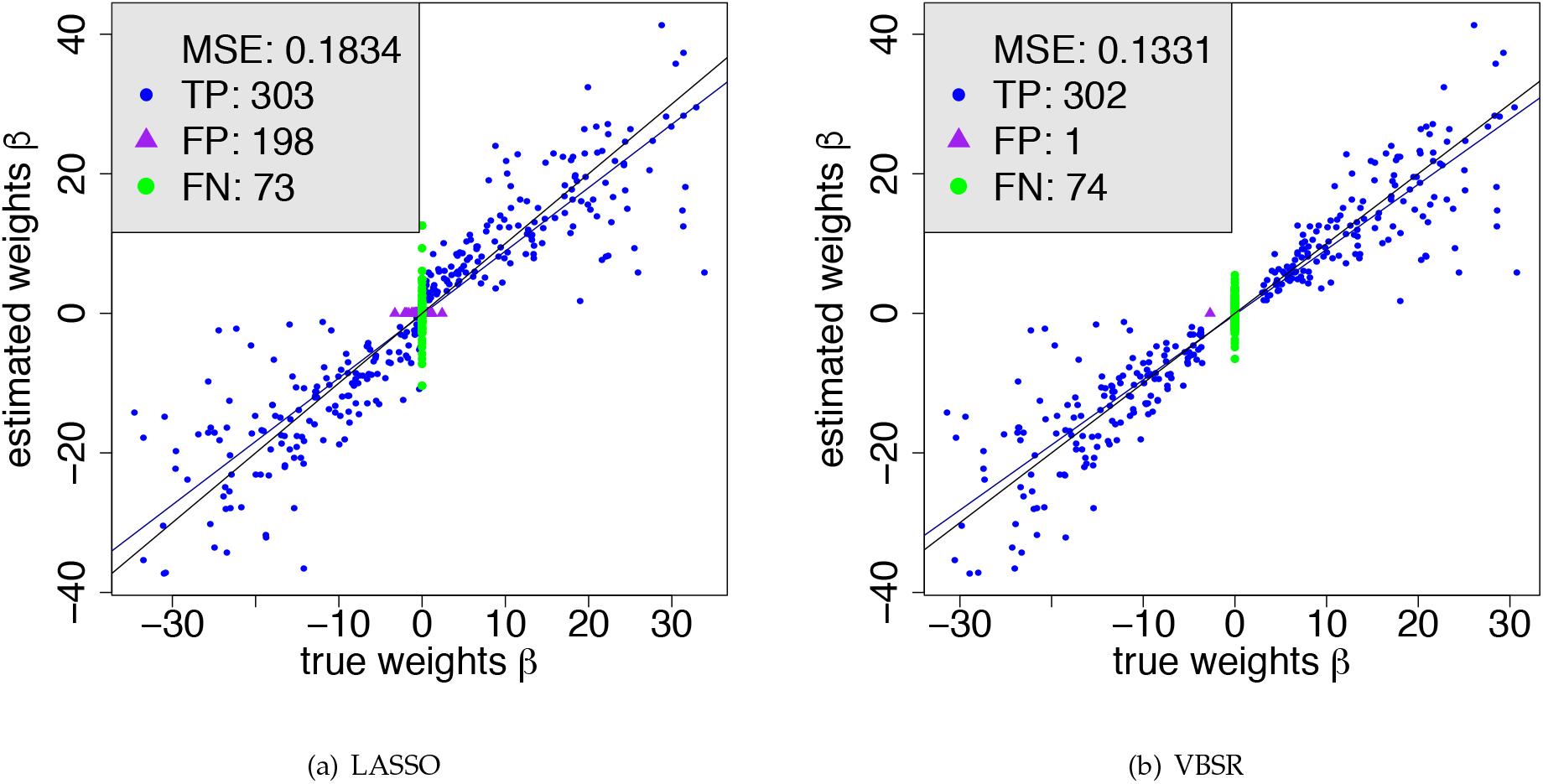
Performance of LASSO (a) and VBSR (b) on estimating the influence of the regulators on the eigengenes of each module when using simulated data and the module assignments are known. Each point on the plots represents the the estimated vs real influence of one regulator across for a given module and bootstrap. For ease of visualization the True Negatives are omitted from the plot.

For the next assessment we simulate the real-case scenario where the module assignments are unknown. We set the number of underlying modules to be *K* = 100 and the expected number of genes per module to be 100, i.e., 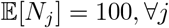. We run the method for module generation described in section* using either LASSO or VBSR.

Since the estimated modules are in principle different from the true ones, we cannot perform the exact same analysis as before. Hence, we first list which regulators are found to have some effect on at least one module and compare this set of regulators with the true ones. If a regulator is found to have an effect (or no effect) in the same number of modules in both the true data and the estimated one, we consider that a true positive (or a true negative for no effect). False positives are those regulators with excess effect found on the estimated modules while false negative are defined as lack of effect.

Figure 4 shows the performance of the proposed method in terms of True Positive Rate (TPR) vs False Positive Rate (FPR) (left) and Precision vs Recall (right) when using either LASSO or VBSR. Note that TPR and Recall are equivalent, but for consistency with the literature we name it differently when pairing it with FPR or Precision. Each black point in the figure represents the performance for a single bootstrap. The performance when the cluster assignments are known is also shown in the figures in red.

**Figure 4:**
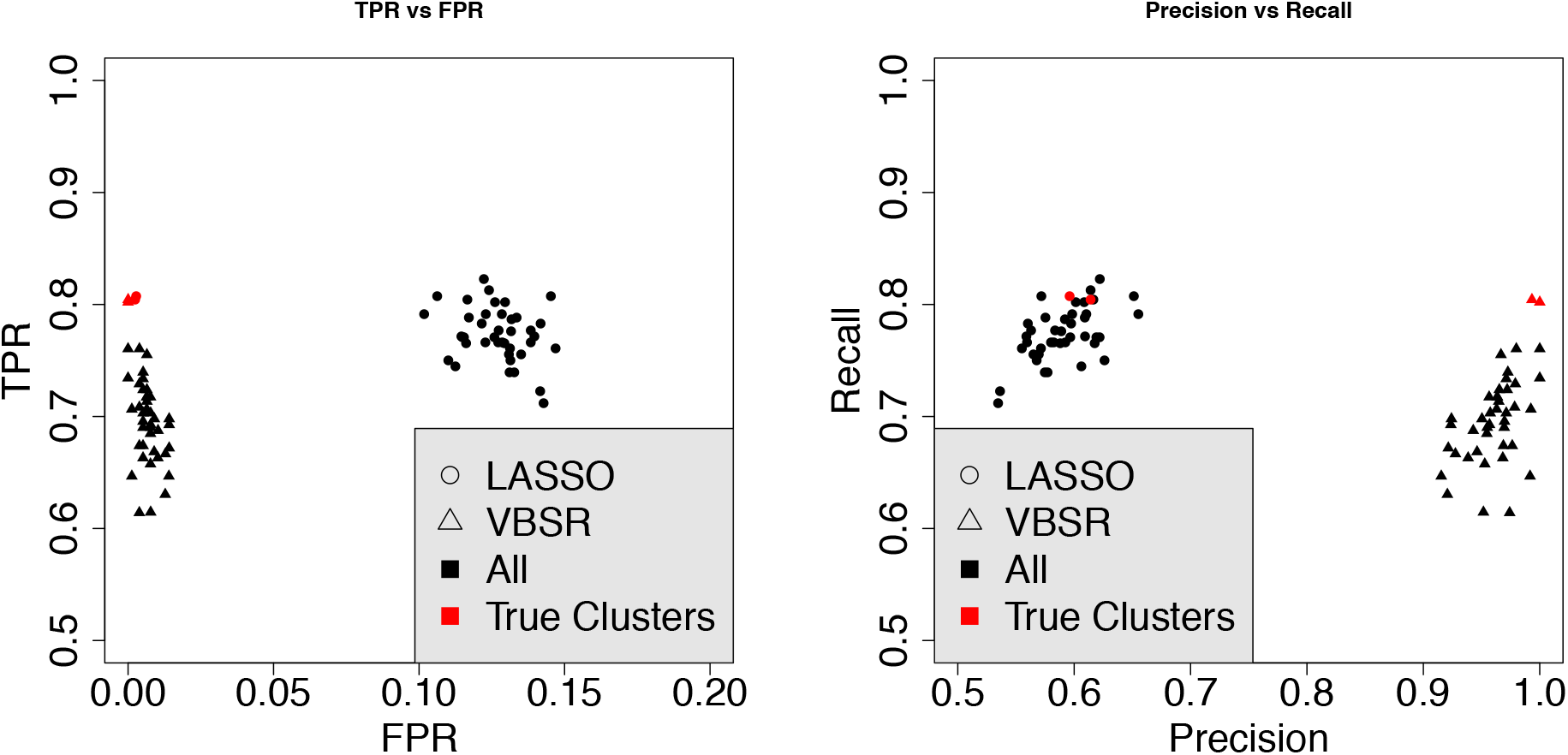
Performance of the estimation of the effect of the regulator on the eigengenes when using either LASSO or VBSR in terms of False Positive Rate vs True Positive Rate (left) and Precision vs Sensitivity (right). The performance when the module assignments are inferred by the method is shown in black, the case where the true module assignments are used is shown in red.

Even though LASSO slightly outperforms VBSR in terms of TPR (or Recall), VBSR clearly outperforms LASSO in both Sensitivity and FPR. Hence, VBSR shows a better trade-off performance in both plots than LASSO. Regarding the degradation incurred by not knowing the true module assignment, for VBSR the same overall performance is observed in terms of FPR and Precision while only a slight degradation of performance is observed for TPR (or Recall). For LASSO the opposite occurs, similar performance is observed for Recall (or TPR) and Precision, however significant degradation occurs in terms of FPR. Moreover, even when the module assignments are known VBSR clearly outperforms LASSO in terms of Precision.

Finally, Figure 5 shows the performance of estimating the different 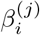 with LASSO (left) and VBSR (right) in the case when the module assignments are unknown. We only show the 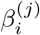 that had a non-zero value in only one of the modules and it was correctly estimated to have a non-zero value by the method. Similar to when the module assignments are known, VBSR (right) outperforms LASSO (left).

**Figure 5:**
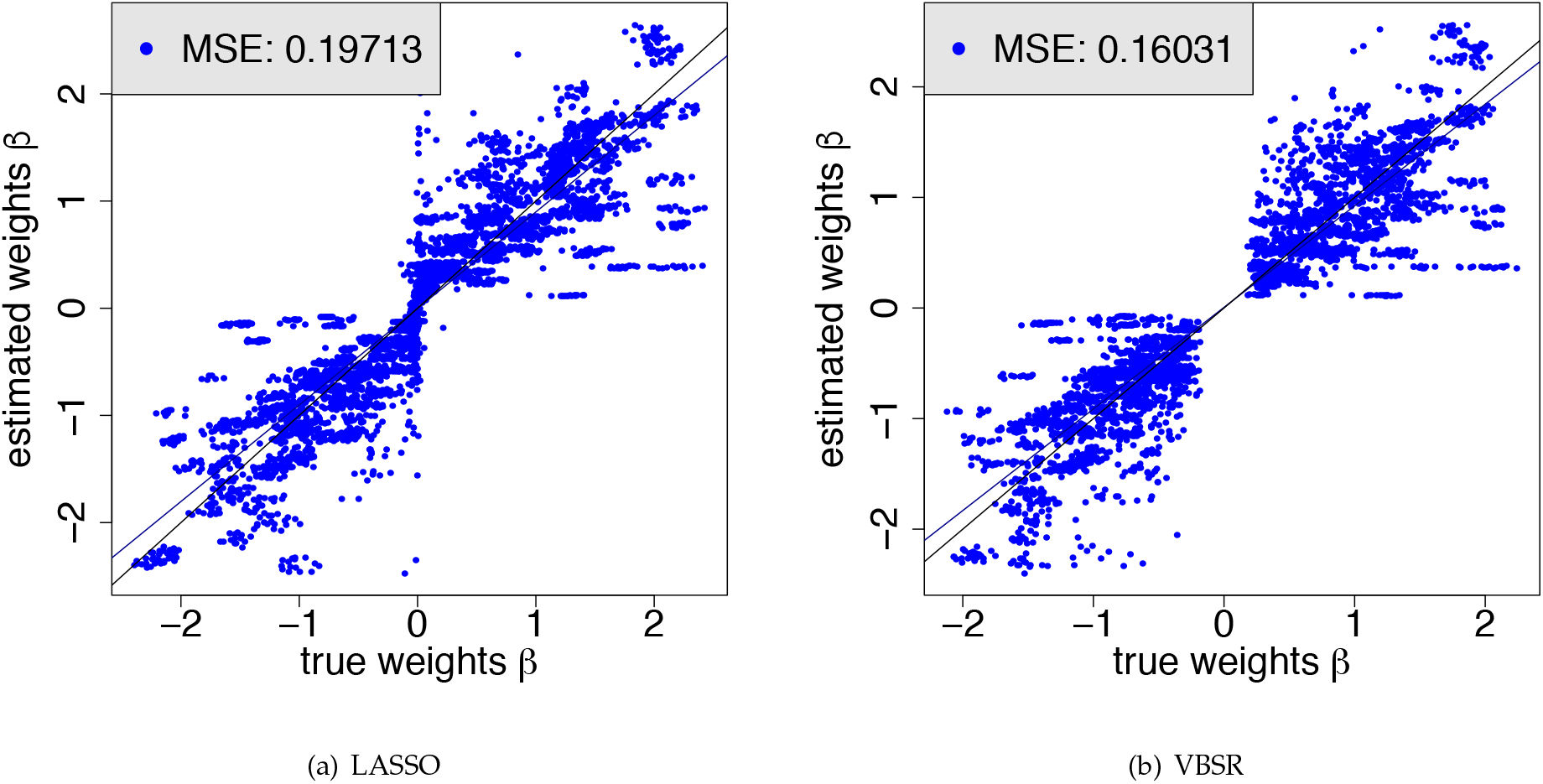
Performance of LASSO (a) and VBSR (b) on estimating the influence of the regulators on the eigengenes of each module when using simulated data and when the module assignments are inferred by the method. Each point on the plots represents the estimated vs real influence of one regulator across a given module and bootstrap. For ease of visualization, only the True Positives are shown.

#### Data cluster analysis

In this section* we focus on assessing the effect that VBSR and LASSO have on the clustering capabilities of the proposed method on simulated data. Since we have knowledge of the true module assignments we use the well-known metrics Variation of Information (VI) (Meilă 2003) and the Adjusted Rand Index (ARI) (Hubert and Arabie 1985) to asses the performance. VI is a proper distance that measures the shared information between the true modules and the modules generated by the method. A VI distance of zero means that the generated module is equal to the truth. On the other hand, the Rand Index is related to how accurately the generated modules represent the true assignments. An adjustment is made to the traditional Rand Index to account for the chance grouping of elements. Hence, an ARI value of one represents perfect accuracy.

Figure 7 shows the clustering performance of the proposed method when using either LASSO or VBSR. As shown in the figure, using LASSO to link regulators with eigengenes yields a better clustering performance for both metrics. We believe that the reason for such improvement is that when using LASSO a cross-validation is performed in order to obtain the regression parameter *λ* that best fits the data. Hence, the heuristics introduced by the cross-validation allow to explore different solutions. Note that VBSR also uses a random initialization of the algorithm, but seems more robust to multiple initializations. More specifically, we observed that for the same eigengene, two runs of LASSO may achieve different solutions (i.e., different value of *β*^(*j*)^), whereas two runs of VBSR seems to consistently achieve the same solution.

**Figure 7:**
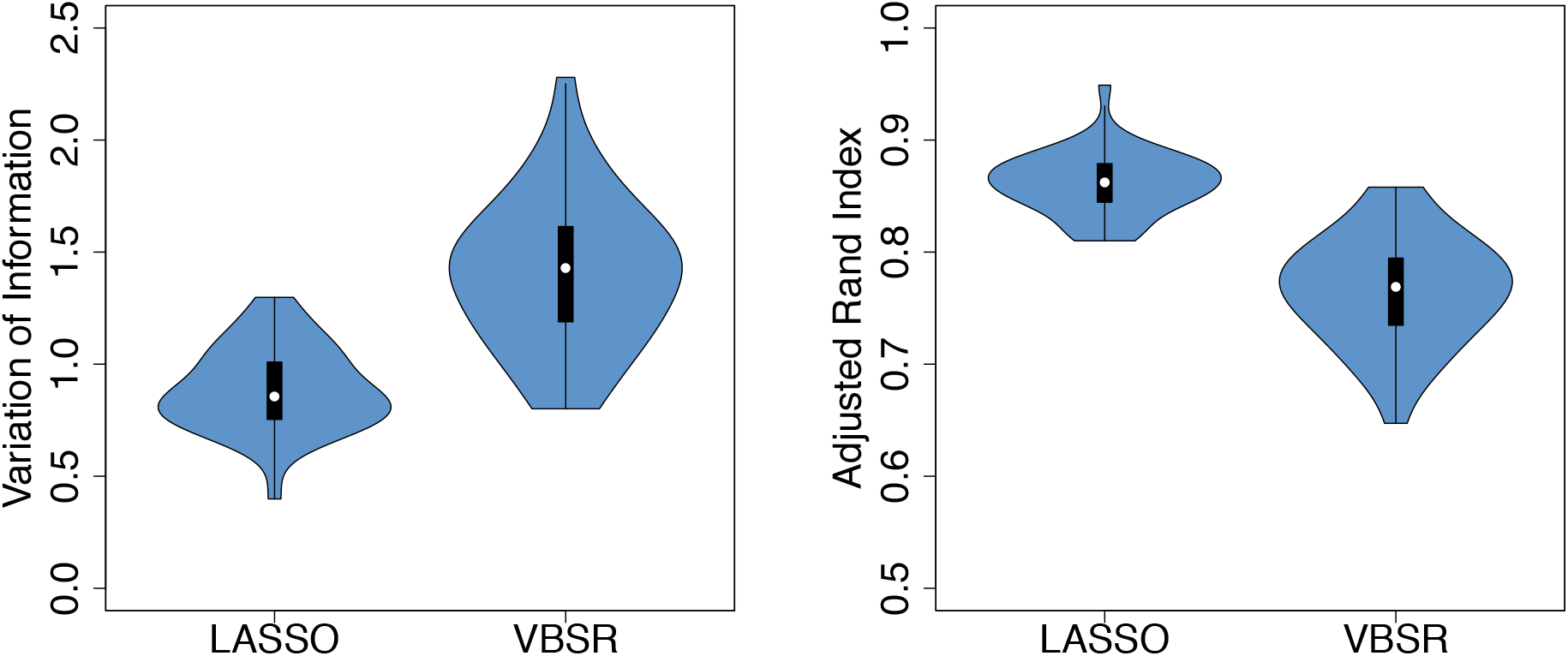
Clustering performance of the proposed method when using either LASSO or VBSR in terms of Variation of Information (left) and Adjusted Rand Index (right).

### Performance analysis on TCGA data

For the next set of experiments, we use the TCGA-OV and TCGA-HNSC datasets, the latter relegated to the Supplementary data.

#### Fitting and clustering performance

First, we focus our assessment on the adjusted R^2^ (also known as coefficient of determination) between the eigengene of a module and its representation using regulators. Ideally, the value of the adjusted R^2^ should be 1 for all modules, as it means that the approximation by regulators perfectly explains all variation across samples occurring in the eigengene.

In addition, to assess the clustering performance when true labels are not available we use the percentage of variance explained by the eigengene as a metric. Since the goal is to generate modules of similarly-regulated genes, the eigengene of modules that are composed of highly-correlated genes will have an eigengene that explains most of the correlation (or variation) existing within the cluster. Whereas if bad assignments are made, the eigengene would only explain a small portion of all the variation. Note that this metric is useful when comparing two methods (relative performance) rather than when measuring the performance of one method (absolute performance) since the true variance in the data is unknown.

Figure 6 shows the performance when using either LASSO (blue) or VBSR (red) to link the expression of the eigengene with the expression of few regulators. Each line represents a bootstrap, the solid lines are the training performance, that is, the performance over the data that was used for computing the modules, and the dotted lines represent the test performance. In order to compute the latter, we perform an 80-20 split of the samples. We use the 80% for training and apply the generated modules to the unused samples. We then compute the adjusted R^2^ between the module’s eigengene and its approximation using regulators (left) as well as the percentage of variance explained by the eigengenes (right).

**Figure 6:**
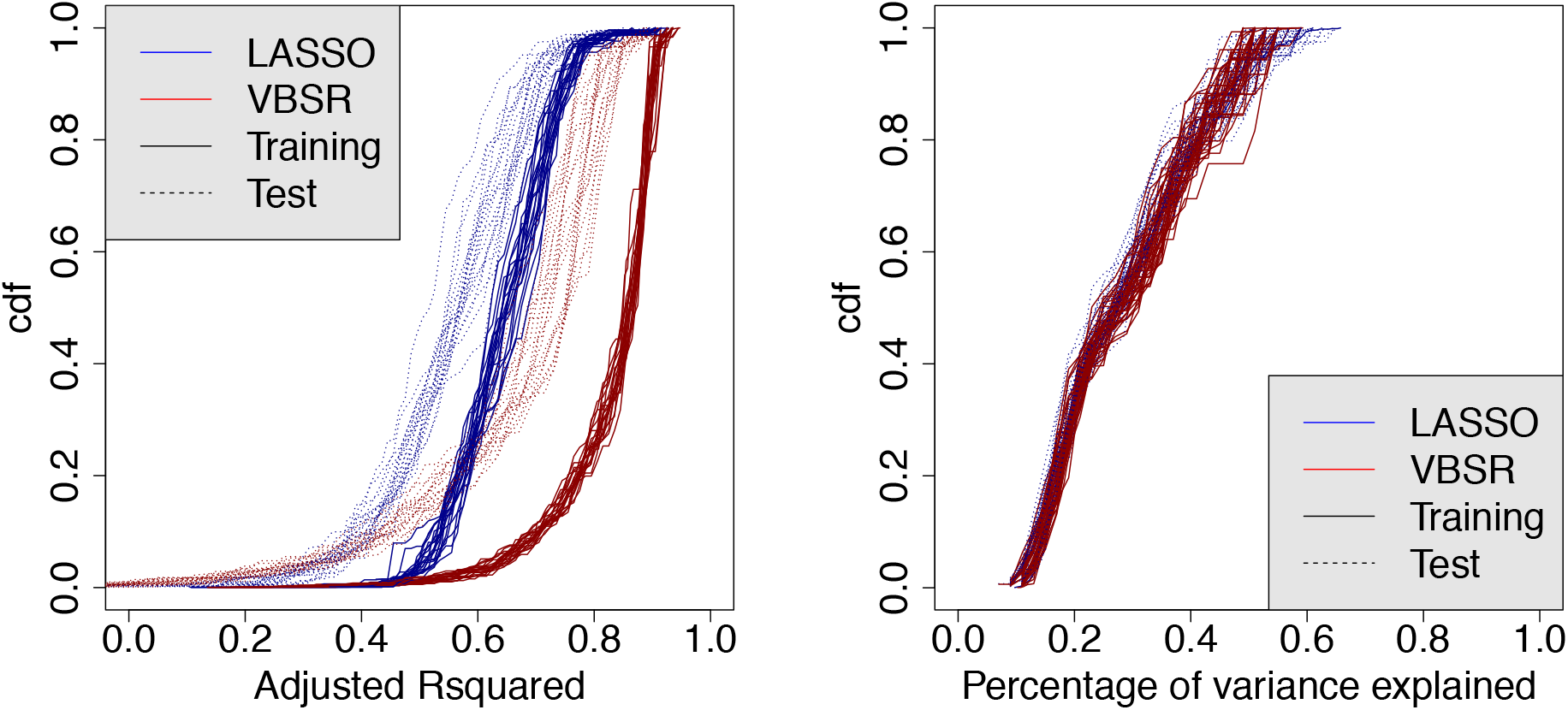
Cumulative Distribution Function (CDF) of the Adjusted R^2^ between the eigengene and its approximation by regulators (left) and the percentage of variance explained by the eigengene (right), for LASSO (blue) and VBSR (red). In solid lines the performance on the training set and in dotted lines the performance on the test set. The CDF is computed across all modules and each curve represents one bootstrap.

These results show that VBSR clearly outperforms LASSO in terms of adjusted R^2^ in both training and test data for all bootstraps; whereas the performance in terms of variance explained by the eigengene is almost identical between the two approaches. This aligns with the results shown for the simulated data where VBSR estimates the regulator-target gene relationships better than LASSO, whereas both perform similarly in terms of clustering capabilities. We observed similar performance for the TCGA-HNSC dataset (results not shown).

In what follows we focus our assessment on analyzing the topology of the resulting networks and the discovery rate of enriched elements. For these analyses we consider all four methods as described in section* ((LASSO_*min*_, LASSO_1*se*_, LM and VBSR)) and use them to either link the module eigengenes with the regulators and/or generating the edges of the bipartite graphs. For comparison, we also consider the cases where each of the methods is used to construct a single bipartite graph directly from the initial expression data.

#### Network Topology Analysis

It has been shown that gene-regulatory networks generally behave as a scale-free network (Barabasi and Oltvai 2004). Figure 8 shows the degree distribution in log-log scale of the proposed method for target genes (a) and regulators (b) when using one of the four proposed methods for linking module eigengenes with the regulators and for generating the edges of the bipartite graphs. The average number of target genes and regulators across the generated bipartite graphs are shown in the corresponding legends as mean±standard deviation. As shown in the figure, VBSR is the sparsest of all the approaches generating an average of 64 target genes and 6 regulators per graph. Also, it is the only method that bounds significantly the degree of target genes linking them with at most 10 regulators. On the other hand, when using LM the average number of regulators per graph increases to 134 (with some including more than 300), while the average number of target genes is 71 with some graphs containing more than 100. Both LASSO approaches lie in between VBSR and LM, with LASSO_*min*_ having, like the LM approach, on average more regulators than target genes per bipartite graph. Hence, the solutions generated by both the LM and LASSO_*min*_ approaches are, in principle, undesirable since their modules (and consequently the associated bipartite graphs) contain, on average, more regulators than target genes.

**Figure 8:**
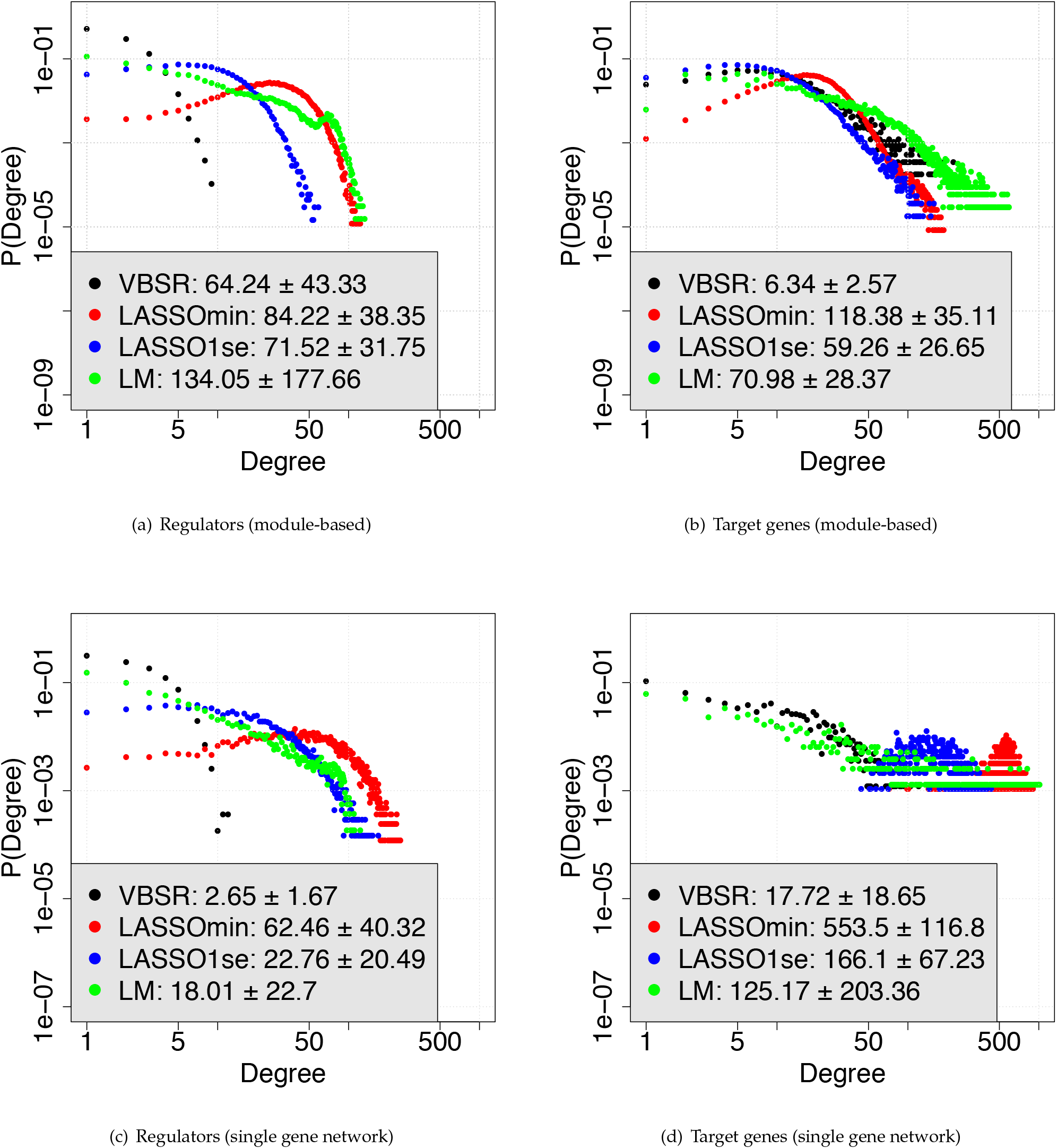
Degree distribution of the bipartite graphs for the regulators (a) and target genes (b) when using the module-based approach. The average number of regulators and target genes for each linking method is shown in the legend. Figures (c) and (d) show the degree distribution for regulators and target genes, respectively, for the case in which a single gene network is generated directly from the expression data. In this case the number in the legend shows the average degree of the regulators and target genes.

Regarding the scale-free topology of the networks, the ones generated using either VBSR or LM are closer to have a scale-free topology than those generated using either of the LASSO approaches.

Figure 8 also shows the degree distribution of the networks generated when either of VBSR, LASSO_*min*_, LASSO_1*se*_ or LM is directly applied to the initial expression data to generate a single gene network. In this case the numbers in the legends show the average degree of the regulators and target genes, respectively as mean±standard deviation. As shown in the figure, both LASSOs generate random networks rather than scale-free ones. Moreover, the average degree distribution of regulators and target genes for both LASSOs is significantly larger than for VBSR and LM. VBSR generates again the sparsest of the networks with an average regulator and target gene degree of 17 and 3, respectively.

Let us now consider the behavioral differences between the module-based and the single gene network approach. Note that this analysis is a proxy for when considering different number of initial modules to discover, which is not shown here due to space constrains. As we reduce the number of modules, which is equivalent to increasing the search space across regulators, both LASSOs generate networks closer to random networks, which is undesirable. For LM, the average sparsity of the networks drastically changes, with more dense networks on the module-based approach. This difference in behavior arises from the fact that the Bonferroni correction used in LM to set the threshold for a meaningful regulator-target gene relationship is more stringent as we increase the search space for relevant regulators. On the other hand, VBSR generates similar networks in both cases, however, in the single gene network approach the sparsity becomes too large to be desirable (most of the regulators are either not present or with degree less than 3).

The same trend of results was observed when different combination of methods were used for linking eigengenes and for generating the edges of the bipartite graphs (Supplementary Figure S1 and S2). Furthermore, when the TCGA-HNSC dataset was used, the results showed also the same trends (Supplementary Figures S3-S5).

In summary, using VBSR to create the modules in the module-based approach or using the proposed LM method to create a single gene network using the initial expression data seems to generate the most desirable networks. Moreover, once the modules are created using VBSR, the choice of method to create the links in the bipartite graph does not significantly affect the topology of the networks (Supplementary Figures S1-S4).

#### Gene set Enrichment Analysis

In order to analyze the biological relevance of the networks generated by the different methods we perform the following analysis: for every regulator in the network (i.e., set of bipartite graphs) we analyze if the target genes connected to it are enriched in curated gene sets. Specifically, we use gene sets from the following databases: BIOCARTA, REACTOME (Fabregat et al. 2015), KEGG (Kanehisa et al. 2016) and GENESIGDB(Culhane et al. 2011).

For the enrichment analysis we use the Fisher’s exact test and we correct for multiple testing by setting the False Discovery Rate (FDR) to 0.05. Furthermore, since the different methods generate networks containing different sets of regulators, we further correct each of the obtained p-values by applying a Bonferroni correction across all the regulators appearing on the corresponding set of bipartite graphs.

Throughout the different bipartite graphs generated across bootstraps there will be regulators that appear multiple times and each time its associated target genes may be enriched in different gene sets. Hence, in what follows, we analyze the following sets of enriched elements: GEA_*reg*_ is the set of regulators whose associated target genes are found to be enriched in at least one known gene set; GEA_*set*_ is the set of known gene sets that are found to be enriched in at least one group of associated target genes; and GEA_*regset*_ is the set of regulator-gene set pairs that are found to be enriched in at least one set of associated target genes.

Let us consider the following example with two regulators *r*_1_ and *r*_2_. As a result of the bootstrapping regulator *r*_1_ is associated to two different set of target genes 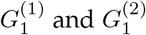 and regulator *r*_2_ is only associated to one set of target genes 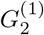. Now, let us assume that both 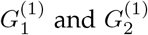 are enriched in the gene set 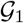, with p-values 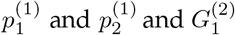 is enriched in the gene set 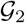 with p-value 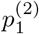.

Then, the above mentioned sets of enriched elements would be as following: GEA_*reg*_ would contain *r*_1_ with p-value 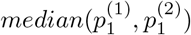 and *r*_2_ with p-value 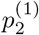; GEA_*set*_ would contain 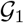 with p-value 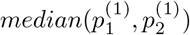 and 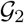 with p-value 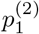; finally, GEA_*regset*_ would contain 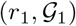 with p-value 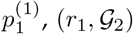 with p-value 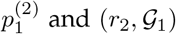 with p-value 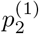.

Finally, due to the vastly different sizes of the generated networks we focus our analysis on enriched elements per network regulator. The reason being is that significantly larger networks will contain more enriched elements mostly due to its size. However, we are generally interested on generating highly informative networks, that is, networks containing a great proportion of enriched elements that are easily interpretable biologically.

Figure 9 shows the cumulative counts per network regulator across the different p-values for each of the above mentioned set of enriched elements, for all gene set databases. The different colors in the figure represent different methods for computing the modules and the different markers on the lines represent the different methods for drawing the edges on the bipartite graphs. The *NET* legend stands for the single gene network approach.

**Figure 9:**
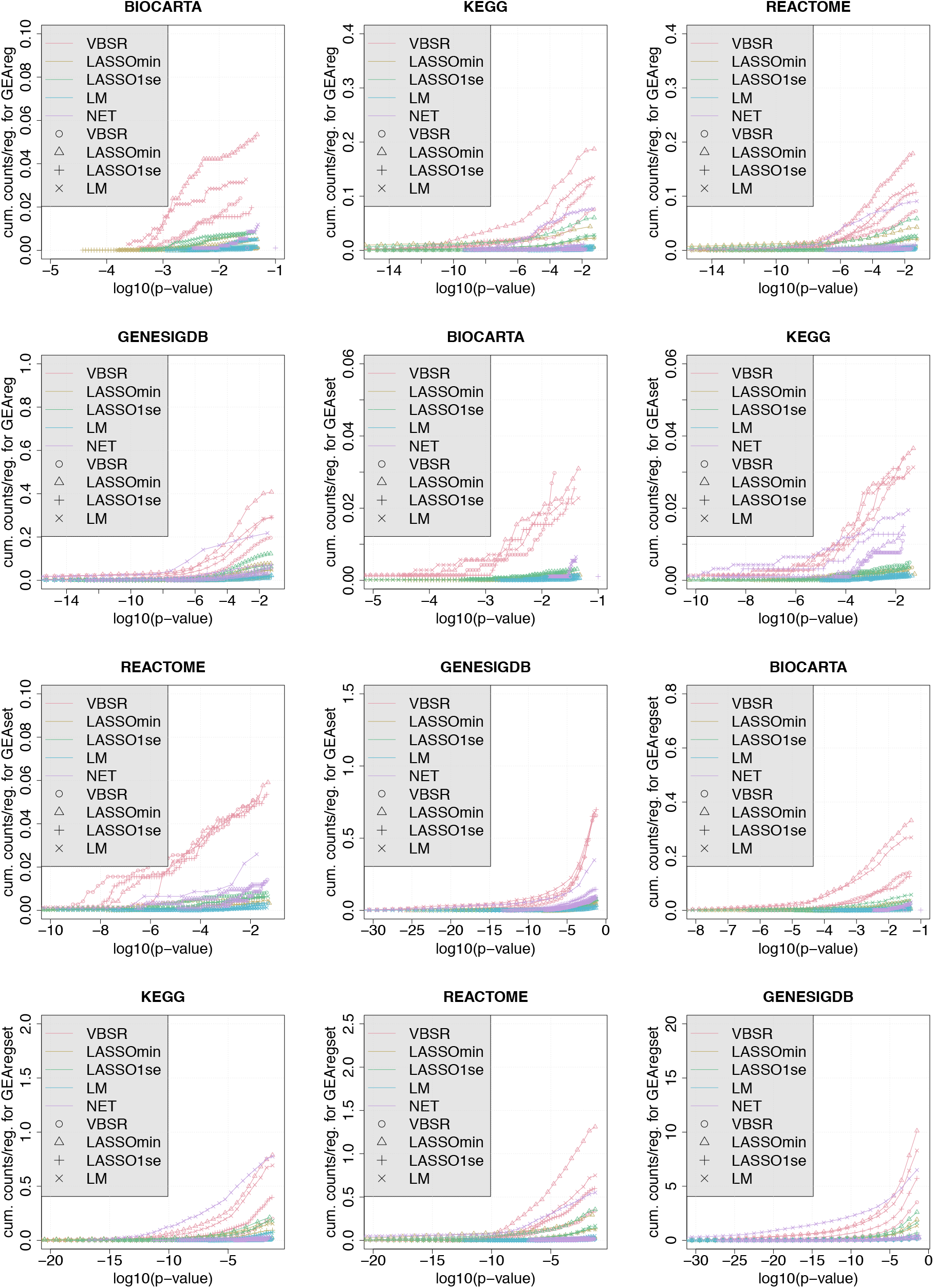
Cumulative counts per network regulator for the three types of enriched elements, GEA_*reg*_, GEA_*set*_ and GEA_*regset*_, for the four analyzed known gene sets. The p-values shown on the x-axes are after the multiple testing corrections. The different colors in the figure represent different methods for computing the modules and the different markers on the lines represent the different methods for drawing the edges on the bipartite graphs. The *NET* legend stands for the single gene network approach.

As seen in the figure, using VBSR to create the modules (pink curves) improves considerably with respect to the other methods for all three types of enriched element sets and for all analyzed gene set databases. In comparison, applying VBSR directly to the initial expression matrix (the single gene network approach, purple curve, circle markers) is among the worst performing of all methods.

The single gene network approach (purple curves) produce, in several cases, less informative results than most of the module-based approaches, with the exception of module-based LM. Interestingly, the only non-module approach that performs, in some cases, similarly to the module-based VBSR is LM. Therefore, while VBSR benefits greatly of the module-based approach, applying LM in a module-base fashion significantly reduces the performance with respect to the non-module version. This is expected as the sparsity of the networks generated by LM changes drastically when considering modules (see previous section*). Again, both LASSOs perform in between VBSR and LM for both the module-based and single gene network approaches.

When analyzing in more detail the module-based VBSR approach (pink curves), we can observe that once the modules are created using VBSR, the choice of method to draw the edges on the bipartite graph makes little difference in some cases but in other cases choosing LASSO_*min*_ (triangle markers) can boost the performance significantly. It is interesting to notice that LASSO_*min*_ is the least sparse solution after LM. This seems to indicate that the module-base approach highly benefits from very sparse solutions to build the modules, but once the modules are built, it is better to rely on less sparse methods to build the bipartite graph, as very sparse solutions may remove true relationships.

Finally, using LASSO_1*se*_ to build the modules in the module-based approach (green curves) outperforms module-based LASSO_*min*_ (yellow curves) and module-based LM (blue curves) in all cases, and the single gene network approaches in most cases (with the exception of the LM case). Similar performance was observed for the TCGA-HNSC dataset (Supplementary Figure S6).

It is important to note that the methods that produce the most informative networks (i.e., module-based VBSR and LM applied to the initial expression matrix) align with those that produced the most desirable networks (scale-free networks with, in case of the module-base approaches, modules with high target gene/regulator ratio).

Finally, we show the benefit of bootstrapping for the different analyzed methods. Figure 10 shows the newly discovered elements per bootstrap from the GEA_*set*_ and GEA_*reg*_ sets for the KEGG database. Note that the rightmost point in the graph shows the total number of elements discovered across all bootstraps.

**Figure 10:**
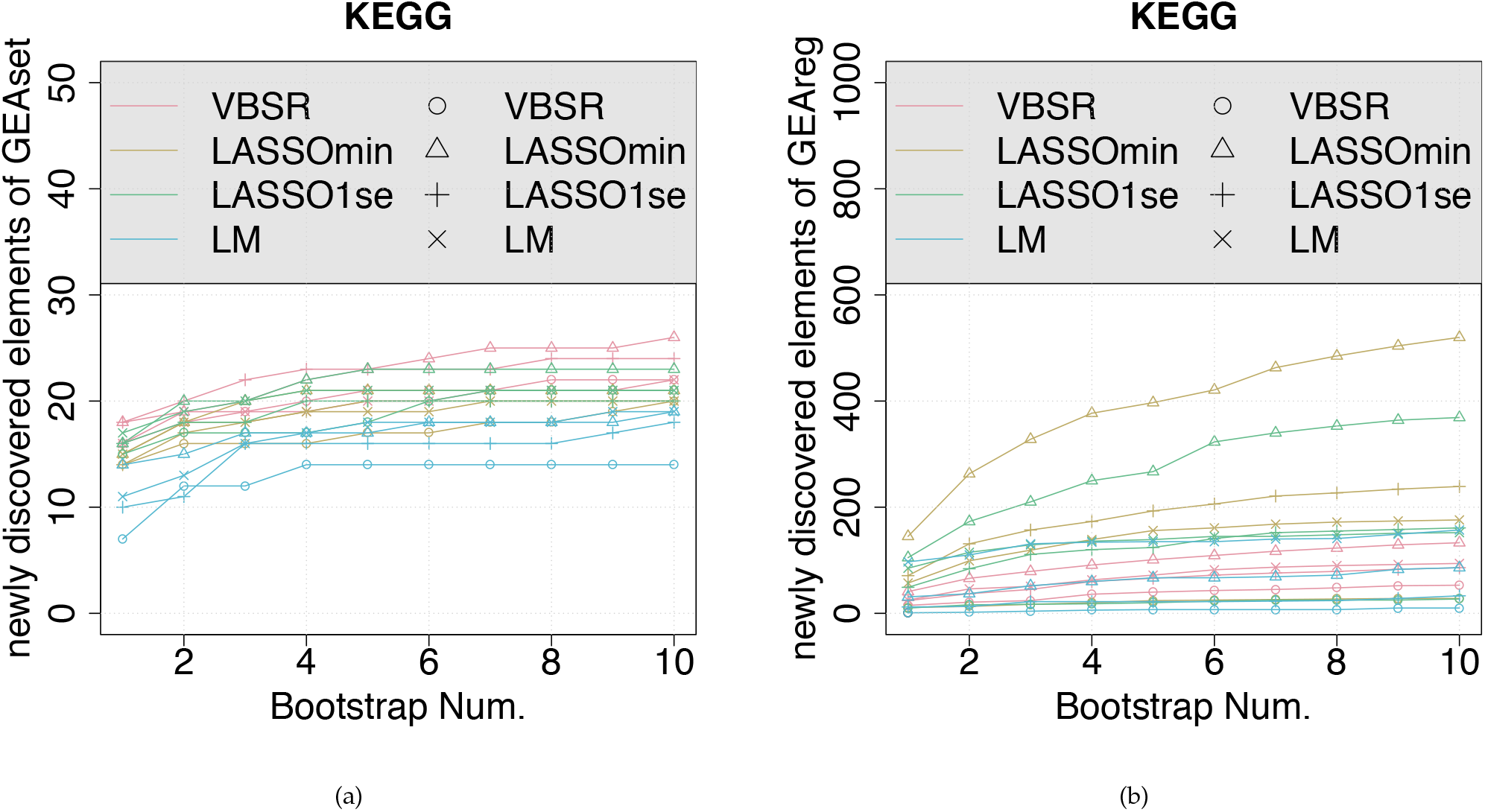
Newly discovered enriched elements from the GEA_*set*_ (a) and GEA_*reg*_ (b) sets across bootstraps when using the KEGG database of known gene sets.

Interestingly, creating modules using VBSR leads to discovering more elements from the GEA_*set*_ set than when using any other method. This is quite significant as VBSR produces the sparsest networks of all methods. On the other hand, for the set of enriched elements GEA_*reg*_, the LASSO approaches (when used either to create the modules or to build the bipartite graphs) lead to the best performance; although, in this case the generated graphs are significantly denser than those generated with VBSR and the topology of the generated networks is not desirable (see Figure 8).

We can also see how certain methods are more robust to bootstrapping, e.g., the LM-LM approach in both figures is almost flat, meaning that after few bootstraps, no more new elements are discovered. This ultimately means that all graphs created across bootstraps are very similar to each other. On the other hand, LASSO results in the largest variation across bootstraps, which aligns with the findings for the clustering performance on simulated data (Figure 7), showing that the cross-validation performed as part of LASSO makes the method less consistent across bootstraps. Finally, VBSR lies in the middle in terms of benefits from bootstrapping.

It is important to note that consistency across bootstrap could be a desirable feature as it may indicate that the method is more robust to over-fitting than methods whose results change significantly across bootstraps. On the other hand, initializing a clustering method with different random initialization settings and choosing the best performing solution is a well established practice when working with clustering methods. In our case we have chosen to focus on the latter, since, as shown in Figure 10 different initialization and sub-sampling of the data yields the discovery of new valid solutions (i.e., new enriched elements that where not discover in previous bootstraps). Moreover, as shown in Figure 6, the proposed module-based VBSR method performs quite well on the test data, reducing overfitting concerns.

## Conclusion

Most of the existing methods for gene regulatory network discovery (also called reverse engineering networks) generate a graph by drawing an edge between two nodes (genes) if their expression satisfies a given constraint. For example, if their correlation is bigger than a given threshold. The remaining methods generally generate a fully connected network where the weight of each edge is computed based on a predefined metric. In this work we show that generating modules of similarly-expressed genes, which are jointly regulated by very few regulators, and then building a bipartite graph on the generated modules, yields more informative networks, as measured by the rate of enriched elements. Moreover, we show that using VBSR as the method to generate the modules rather than LASSO or correlation, which have been the preferred choice in the literature, achieves significantly better results. Finally, we show that bootstrapping is advantageous in this case, mainly due to the heuristic nature of the proposed method. Specifically, it leads to the discovery of new enriched elements that would have been missed otherwise. However, in some cases this also leads to a decrease of the discovery rate of enriched elements as bootstrapping creates larger networks. Nevertheless, when using VBSR to discover the co-regulated modules the use of bootstrapping increases the enriched element discovery rate at a faster pace than the number of new nodes introduced by the bootstrapping.

